## Supplemental Table and Figures for "A multiplexed, automated evolution pipeline enables scalable discovery and characterization of biosensors"

**Table S1 | Selection parameters.**

| Selection | Prefixes | Suffix | Library | Order of Rounds | Ligation Method | Free $Mg^{++}$ | Rounds | Hits (First Round) |
|---|---|---|---|---|---|---|---|---|
| **S1** | A, B | S | Loop1: N7<br>Loop2: N30 | One -ligand selection round for cleavers, followed by alternating +/- ligand rounds. | Separate RT primer, splint, and ligation substrate | Low | 57 | Theophylline (R57) |
| **S2** | A, B, W | X | Loop1: N4-N8<br>Loop2: N30, N60<br>+<br>Loop1: N30, N60<br>Loop2: N4-N8 | Four -ligand selection rounds for cleavers, followed by alternating +/- ligand rounds. | Single molecule RT primer/splint/ligation substrate | +4mM during RT setup after R46 | 64 (T2,T5), 78(T3), 100(T6) | (*S*)-reticuline (R36) |
| **S3** | W, Z | X | Loop1: N4-N8<br>Loop2: N30, N60<br>+<br>Loop1: N30, N60<br>Loop2: N4-N8 | Two -ligand selection rounds for cleavers, followed by alternating +/- ligand rounds. | Single molecule RT primer/splint/ligation substrate | +4 mM during RT | 114 (T1), 202(T2), 198(T3) | (*S*)-reticuline (R126), noscapine (R84), trans-zeatin (R126), aciclovir (R74) |
| **S4** | W, Z | X | Loop1: N4-N8<br>Loop2: N30, N60<br>+<br>Loop1: N30, N60<br>Loop2: N4-N8 | Started with the pre-selection product (R2) from the prior selection (S3), followed by a repeating pattern of one +ligand round followed by two -ligand rounds. | Single molecule RT primer/splint/ligation substrate | Low | 294(T1) | Gardiquimod (R102) |

In above table, N*xx* refers to the number of degenerate nucleotides in each of the ribozyme loops; T*n* refers to ligand group *n*; S*n* refers to selection run *n*; R*n* refers to round *n* of a selection run. Low values for free $Mg^{++}$ are <1mM.

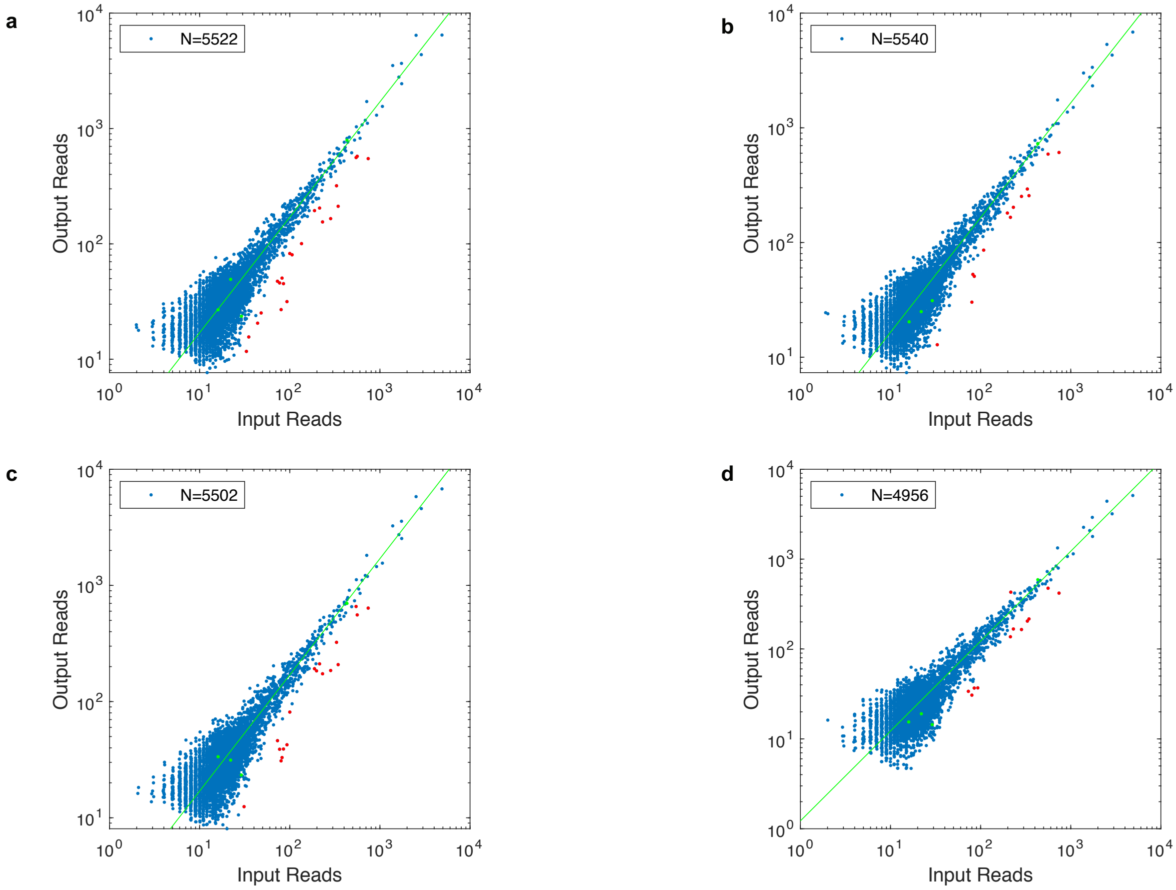


**Fig. S1 | Efficiency of CleaveSeq including regeneration.** CleaveSeq was run on a library of biosensor sequences formed from mixing the products of rounds 91, 98, 106, 114, and 122 of DRIVER select S3 and then constricting to ~5,000 distinct sequences. The total number of reads attributed to cleaved and uncleaved products (each normalized using reference sequences) for each sequence is plotted against the number of reads of the same sequences at the input to CleaveSeq. The green line shows the expected output reads based on the ratio of total NGS reads allocated to the input and output libraries. Red points indicate sequences with output counts significantly lower than other sequences based on a two-proportion z-test with $p<0.01$ and applying a Bonferroni correction for multiple hypotheses. **a.** CleaveSeq with no ligand present, **b.** T1 ligand mixture, **c.** T2e ligand mixture, **d.** T3e ligand mixture.

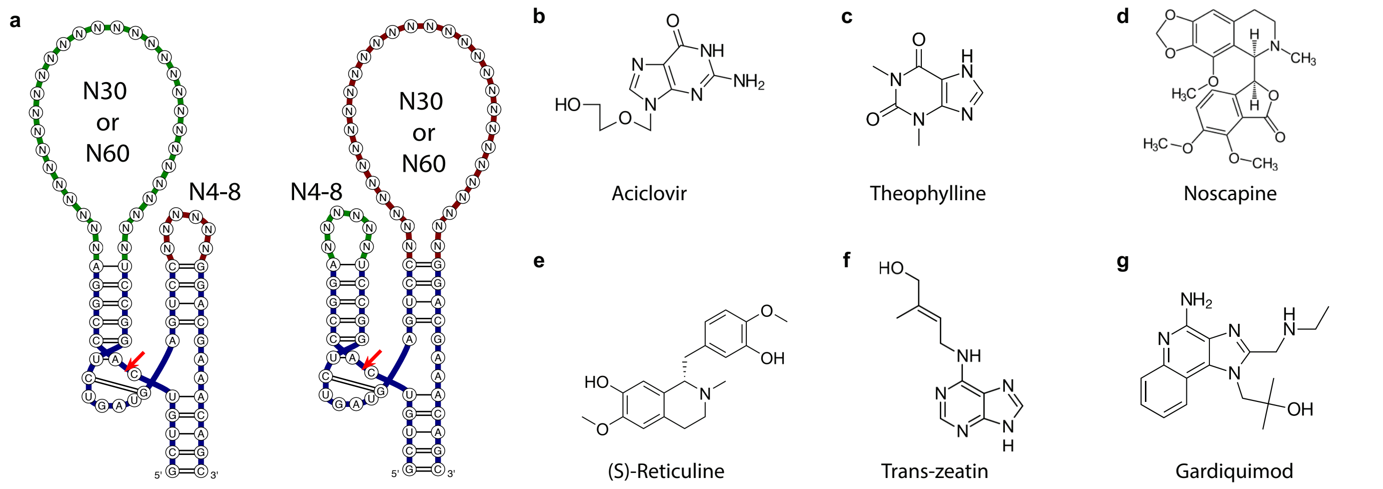


**Fig. S2 | Library design and ligand structures. a.** Secondary structure representation of general biosensor library design with the loop randomizations indicated. N6 small loops and N30 large loops are shown. **b-g.** Chemical structures of ligands for which novel biosensors were validated in this work: aciclovir (**b),** theophylline (**c**), noscapine (**d),** (*S)*-reticuline (**e),** trans-zeatin (**f**), gardiquimod **(g**).

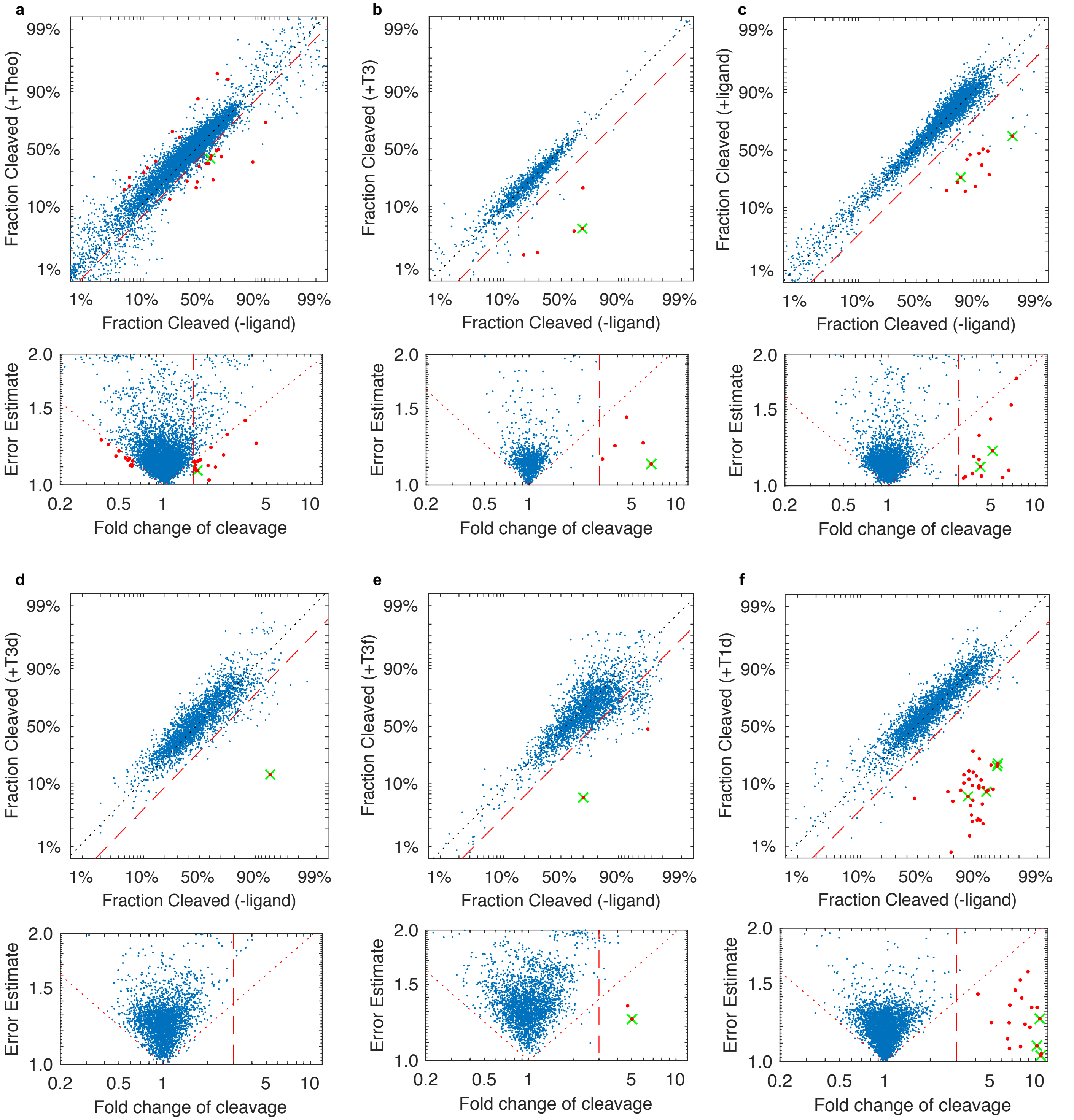


**Fig. S3 | Fraction cleaved and fold change of fraction cleaved for evolved libraries.** Comparison of cleavage fractions in the presence and absence of the ligand mixture determined via CleaveSeq of libraries at various points during the selections. In each section, the top subpanel shows the fraction cleaved of each of the sequences which have at least 30 reads in each of the -ligand and +ligand conditions. Bottom panel, the same data plotted with the ratio of the cleavage fractions in the presence and absence of the ligand mixture on the x-axis and the standard error of the ratio on the y-axis. Dotted diagonal line, delineates the region where a multiple-hypothesis test would reject the null hypothesis of non-switching, with alpha=1/N. In both panels: dashed line, boundary where the fold change of cleavage is at least 3x (or 2x for theophylline); red dots, indicate sequences with strong, significant (i.e., below the diagonal line and to the right of the dashed line) switching; green crosses, indicate validated biosensors that were first identified from the particular analysis. **a.** S1 round 57 against theophylline (Theo-421 in green). **b.** S2 round 36 against T3 ligand group (SRet-584 in green). **c.** S3 round 74 against T1b ligand group (Acic-711 and Acic-758 in green). **d.** S3 round 84 against T3d ligand group (Nosc-786 in green). **e.** S3 round 150 against T3f ligand group (TZea-927 in green). **f.** S4 round 102 against T1d ligand group (Gard-337, Gard-910, Gard-544, and Gard-674 in green).

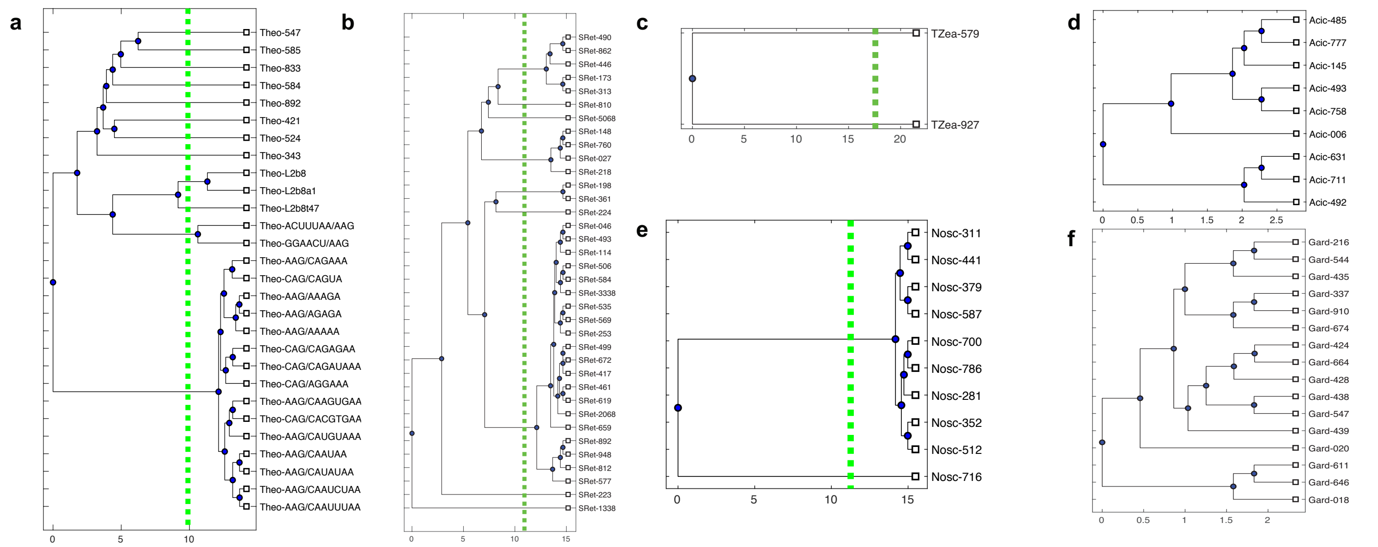


**Fig. S4 | Dendrograms of validated biosensors.** Dendrograms of biosensors found for each distinct ligand based on pairwise distance (number of mismatches) with average linkage. A distance of 5 mismatches (green dotted lines) was used to classify groups of sequences into different families. Dendrograms are provided as follows: **a.** theophylline (8 distinct families found during selection S1). The bottom 20 sensors were previously designed biosensors based on the original TCT8-3 theophylline aptamer), **b.** (S)-reticuline (9 distinct families), **c.** trans-zeatin (2 distinct families), **d.** aciclovir (1 family), **e.** noscapine (2 distinct families), **f.** gardiquimod (1 family).

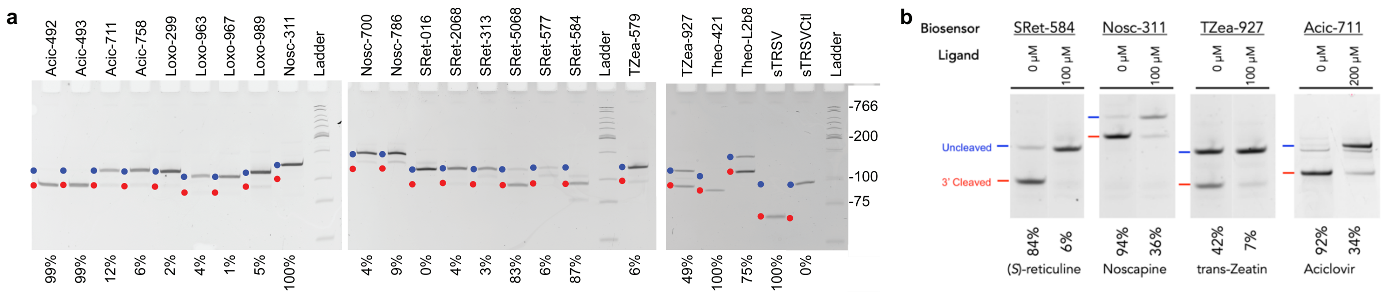


**Fig. S5 | Polyacrylamide gel electrophoresis analysis of RNA biosensors. a.** PAGE gel and analysis of biosensors and controls used to determine the fraction cleaved following a co-transcriptional cleavage assay in the absence of ligand. The labels above indicate the particular biosensor or control RNA sequence (Supplementary Table 3), blue and red dots indicate the expected position of the uncleaved and 3'-cleaved products, respectively; numbers below are the ratio of the cleaved band intensity to the total of the cleaved and 3'-uncleaved bands. **b.** PAGE assay of select biosensors showing fraction cleaved in the -ligand and +ligand conditions.

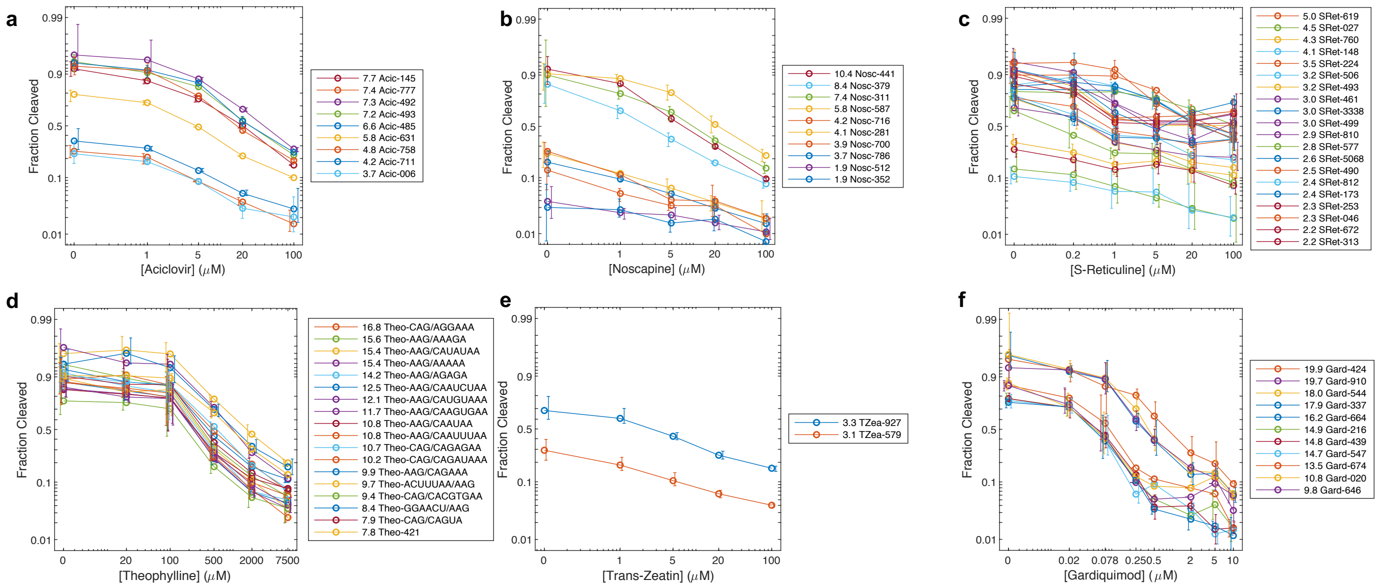


**Fig. S6 | Fraction cleaved for select biosensors as a function of ligand concentration. a-f.** The fraction cleaved of DRIVER-selected biosensors was measured over a range of ligand concentrations using the CleaveSeq assay. Each plot shows the biosensors that exhibit a fold change of fraction cleaved of at least 2.0. Points shows measurements derived from at least 100 NGS reads. Legends include the fold change of fraction cleaved of the indicated biosensor over the ligand range tested. Error bars are [25%,75%] confidence intervals of the mean, calculated over at least 3 replicates; note that error bars are slightly offset from data points to improve legibility. Data is shown for sensors responsive to aciclovir (**a**), noscapine (**b**), (*S*)-reticuline (**c**), theophylline (**d**), trans-zeatin (**e**), and gardiquimod (**f**).

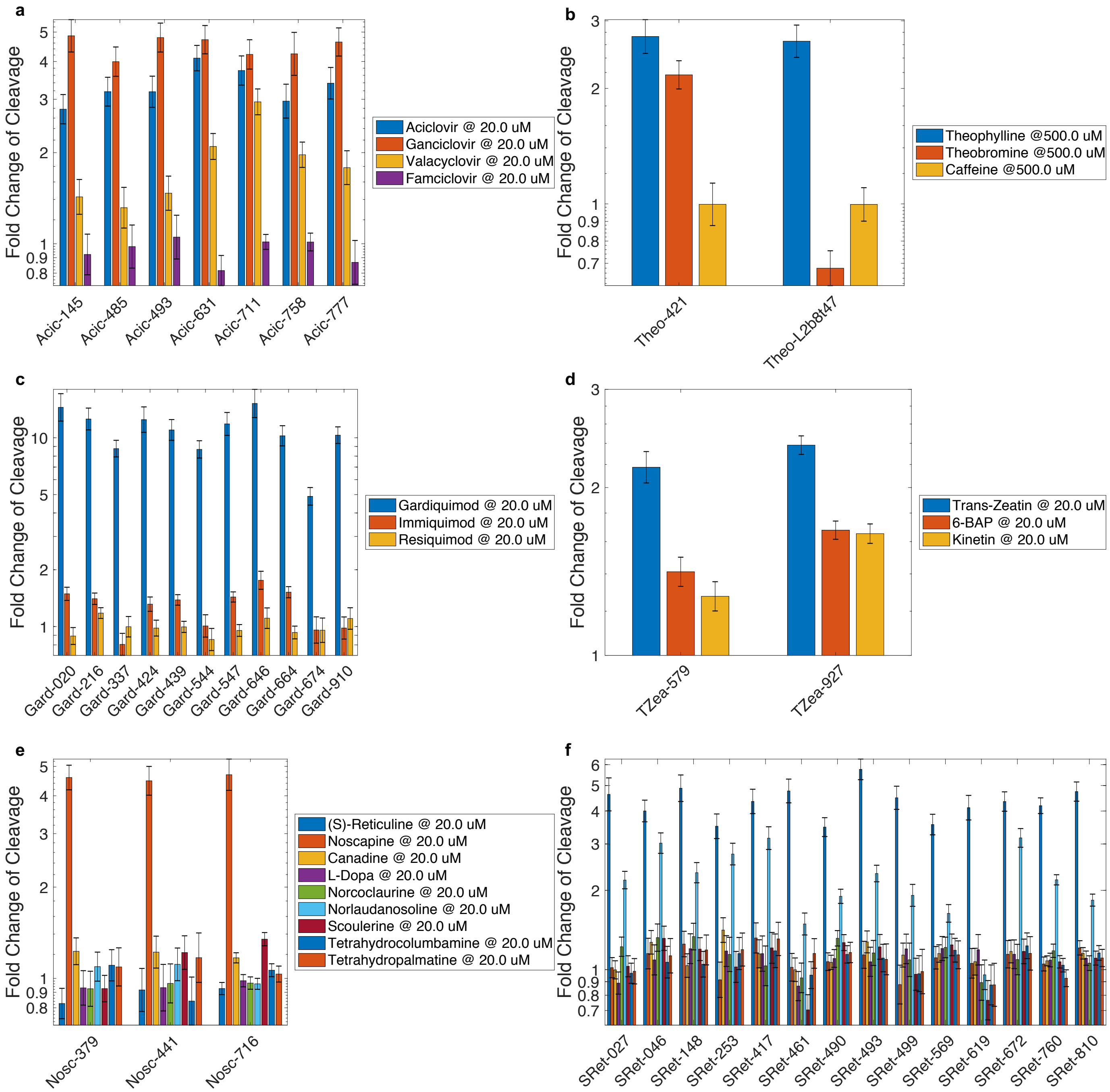


**Fig. S7 | Selectivity of DRIVER Biosensors. a-f.** CleaveSeq was used to measure fold change of cleavage of representative biosensors in the presence of their designated targets and several similar molecules relative to the cleavage in the absence of target. For each condition error bars show the standard deviation of at least two independent replicates. **a.** Aciclovir sensors against aciclovir, ganciclovir, valacyclovir, and famciclovir. **b.** Theophylline sensors against theophylline, theobromine, and caffeine. Theo-L2b8t47 is a previously published sensor based on the TCT8-4 aptamer. **c.** Gardiquimod sensors against gardiquimod, imiquimod, and resiquimod. **d.** Trans-zeatin sensor against trans-zeatin, 6-benzylaminopurine, and kinetin. **e.** Noscapine sensors against several benzylisoquinoline alkaloids (BIAs) and precursors to BIAs: (*S*)-reticuline, norlaudanosoline, noscapine, canadine, L-dopa, norcoclaurine, scoulerine, tetrahydrocolumbamine, and tetrahydropalmatine. **f.** (*S*)-reticuline sensors against the same set of BIAs and precursor BIAs.

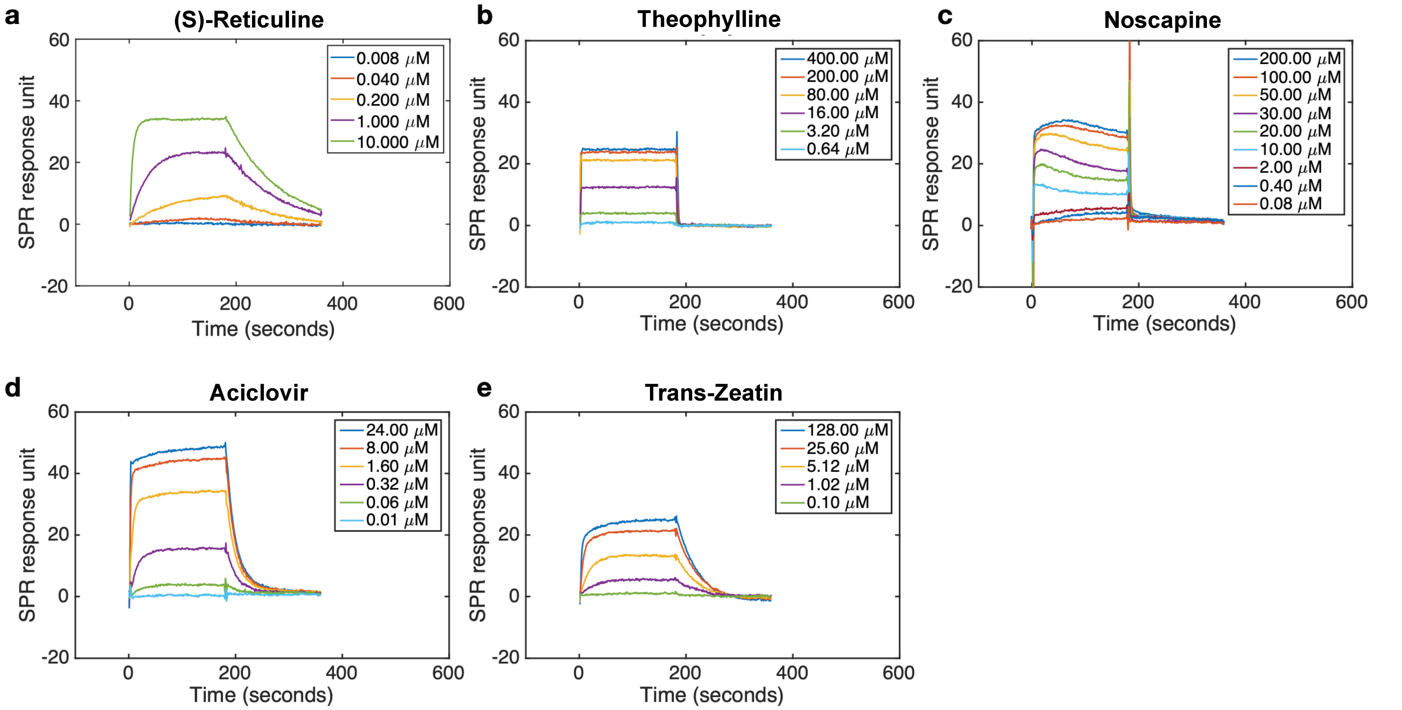


**Fig. S8 | Surface plasmon resonance (SPR) assay sensorgrams to measure the binding affinity of representative biosensors. a.-e.** Select biosensors characterized using Surface Plasmon Resonance from a single Biacore run. Representative sensorgrams are provided for the following biosensors: (S)-reticuline biosensor SRet-584 (**a**), theophylline biosensor Theo-421 (**b**), noscapine biosensor Nosc-786 (**c**), aciclovir biosensor Acic-145 (**d**), and trans-zeatin biosensor TZea-579 (**e**).

a

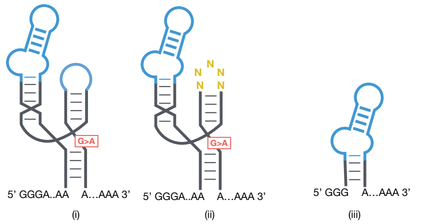


b

| Sensor | $K_D$ (μM) | | |
|---|---|---|---|
| | (i) | (ii) | (iii) |
| **Acic-145** | 1.5 ± 0.3 | 0.680 ± 0.004 | 0.700 ± 0.004 |
| **Nosc-441** | 13 ± 2 | 20 ± 1 | 19.7 ± 7 |
| **SRet-760** | 0.15 ± 0.01 | NB | NB |
| **Theo-421** | 68 ± 20 | 19 ± 4 | NB/17.0 ± 0.7* |
| **TZea-579** | 3.1 ± 0.5 | 5.6 ± 0.4 | 37 ± 3 |
| **TZea-927** | 62 ± 20 | NB | NB |

c

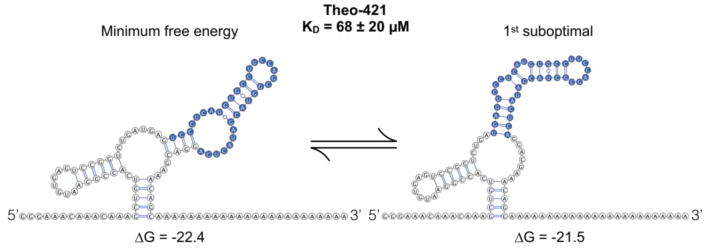


d

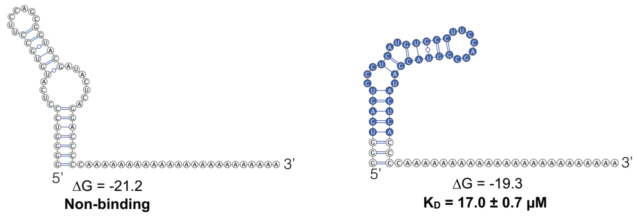


**Fig. S9 | SPR binding affinities of select biosensors and characterization of putative binding domains. a**. Schematic showing the modifications made to the sequences of biosensors used in surface plasmon resonance (SPR) binding characterizations; (i) G12A mutation to prevent ribozyme self-cleavage during binding affinity experiments, (ii) randomizing the smaller loop of the biosensor sequence in addition to a G12A mutation, (iii) truncation to include only the stem I or stem II sequences, whichever contains the larger loop, excluding the catalytic core and the rest of the ribozyme context. **b.** Table shows the equilibrium dissociation constants ($K_D$) as a measure of binding affinity for 6 representative biosensors from five ligand classes (aciclovir, noscapine, (*S*)-reticuline, theophylline, and trans-zeatin) of DRIVER-selected biosensors in the three architectural contexts as illustrated in **a**. Values are mean ± s.e.m of three or more replicate samples. NB, no observed binding. NA, not available. *Theo-421 stem-only (iii) version did not exhibit binding, even though all sequences with preserved binding in (ii) showed binding in (iii); however, the stem-only sequence that stabilizes the first suboptimal structure did, as shown **d**. **c**. Predicted secondary structures of Theo-421 show the minimum free energy structure (top left) and a suboptimal structure (top right), with the predicted free energies ΔG indicated below each structure using the secondary structure folding program RNAstructure. **d.** The truncated structures derived from each of the two structural conformations are shown. $K_D$ corresponds to that in **b**(iii), annotating the structures for clarity. Blue, sequence bases in the putative binding domain.

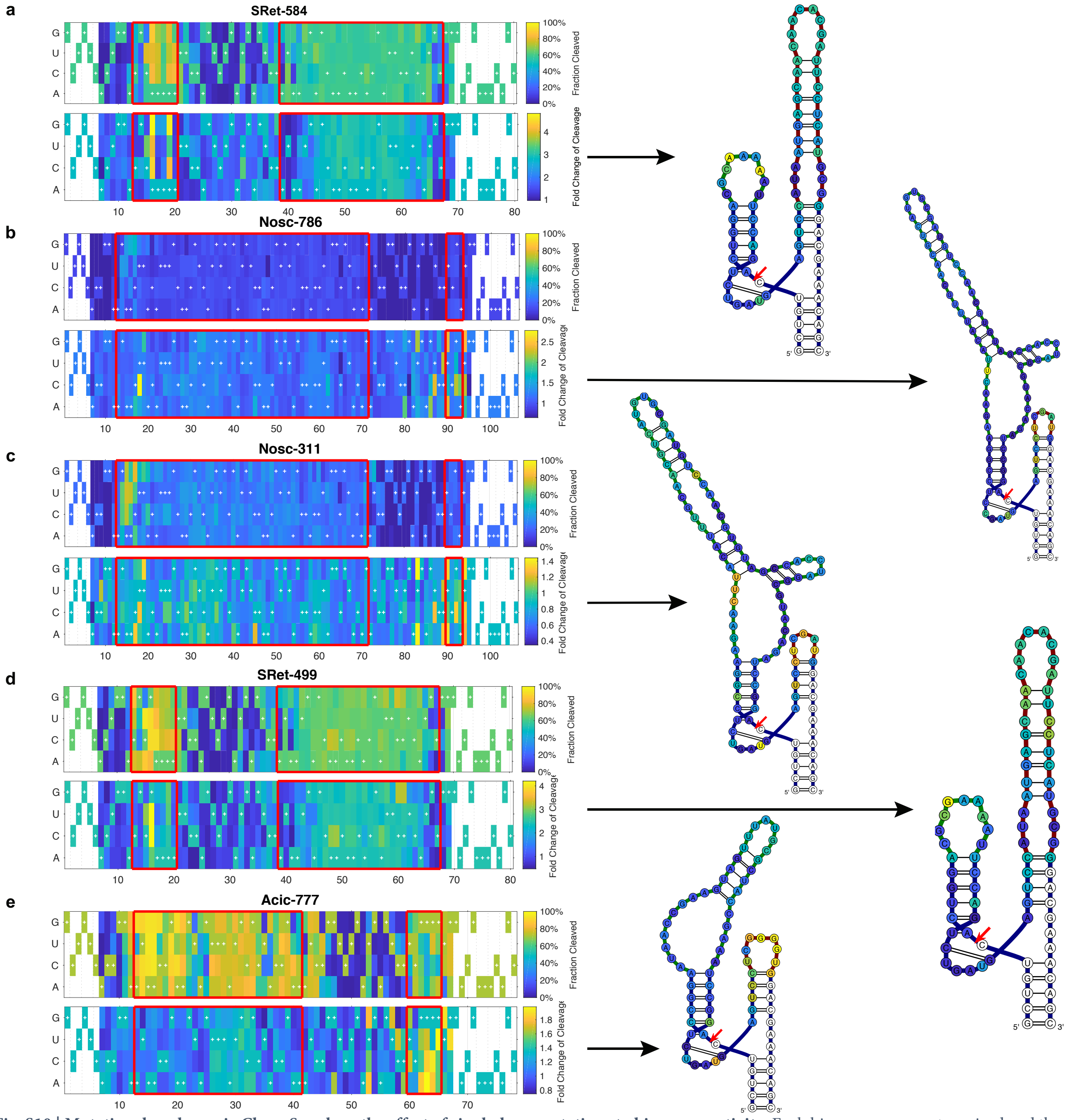


**Fig. S10 | Mutational analyses via CleaveSeq show the effect of single-base mutations to biosensor activity.** Each biosensor was mutagenized and the resulting library characterized via CleaveSeq in the presence and absence of ligand. The top left subpanel of each section shows the fraction cleaved as a function of base identity and position in the biosensor without ligand present. Red boxes delineate the loop I and loop II regions of the biosensor; plus symbols indicate the sequence of the wild-type biosensor. The white regions have no mutation data as they overlapped with primers during subsequent PCR. The lower left subpanels show the fold change of fraction cleaved in the presence and absence of the ligand as a function of base identity and position of point mutations in the biosensor. To the right of each heat map, the fold change of cleavage of the most favorable mutation at each nucleotide position was mapped to the secondary structure of the unmutated biosensor. The color mapping is the same as for the corresponding lower left subpanel. **a.** SRet-584 with (*S*)-reticuline at 200 nM, **b.** Nosc-786 with noscapine at 20 µM, **c.** Nosc-311 with noscapine at 5 µM, **d.** SRet-499 with (*S*)-reticuline at 5 µM, **e.** Acic-777 with aciclovir at 5 µM.

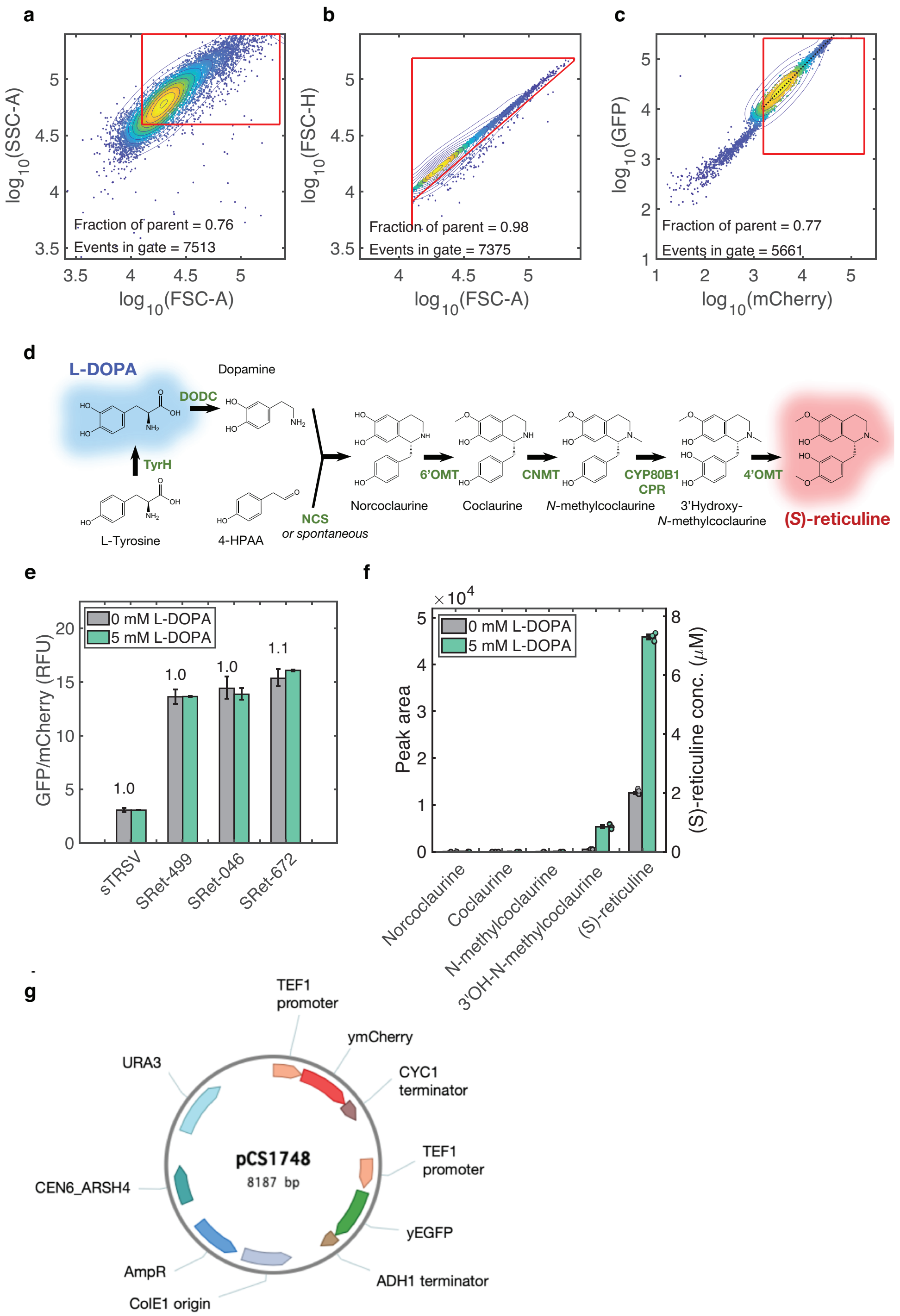


**Fig. S11 | Validation of the *in vivo* gene-regulatory activity of DRIVER-selected biosensors. a.** Representative flow cytometry plot of forward scatter area against side scatter area for an aciclovir switch (Acic-493) assayed for gene-regulatory activity in yeast. Each flow cytometry event is shown as a dot. Viable cells are gated with red margins. Viable cells from **a** are shown on the plot of forward scatter height against forward scatter area. **b.** Singlet cells are gated with red margins. **c.** Viable and single cells from **a** and **b** are plotted with GFP against mCherry fluorescence intensities. **d**. Schematic of a heterologous biosynthetic pathway converting L-DOPA, the fed substrate, to (*S*)-reticuline in engineered yeast strain (CSY1171) used to assay the gene-regulatory activities of (*S*)-reticuline biosensors. **e.** Flow cytometry results of (*S*)-reticuline switches assayed in the yeast strain W303α, which does not express the (S)-reticuline biosynthetic pathway, in 0 mM and 5 mM of L-DOPA. sTRSV, wild-type constitutively active ribozyme. Error bars are standard error of mean of at least 4 biological replicates. **f.** Bar plot of the abundance of BIA pathway intermediates produced in the media from the yeast strain CSY1171 with and without L-DOPA feeding. Peak area refers to the integrated area of the peak detected for each compound by LC-MS/MS using the MRM transitions reported in Materials and Methods. Error bars indicate s.e.m; individual filled circles correspond to biological replicates. Absolute concentration quantities of (S)-reticuline only was obtained from fitting to a standard curve and are indicated on the right vertical axis.**g**. Plasmid map of pCS1748, a dual color fluorescent reporter plasmid used in characterizing ribozyme switch-based biosensors in yeast cells. Biosensor sequences are inserted in the 3' untranslated region via cloning, after the stop codon of yEGFP and before the start of the ADH1 terminator.
